# Colloblasts act as a biomechanical sensor for suitable prey in *Pleurobrachia*

**DOI:** 10.1101/2020.06.27.175059

**Authors:** JP Townsend, GOT Merces, GP Castellanos, M Pickering

## Abstract

Ctenophores are a group of largely-planktonic, gelatinous carnivores whose most common method of prey capture is nearly a phylum-defining trait. Tentaculate ctenophores release an unknown proteinaceous adhesive from specialised colloblast cells lining their tentacles following prey contact with the tentacles. There exist no extant studies of the mechanical properties of colloblast adhesive. We use live microscopy techniques to visualise adhesion events between *Pleurobrachia pileus* colloblasts and probes of different surface chemistries in response to probing with varying contact areas. We further define two mechanisms of adhesion termination upon probe retraction. Adapting a technique for measuring surface tension, we examine the adhesive strength of tentacles in the ctenophore *Pleurobrachia bachei* under varying pH and bonding time conditions, and demonstrate the destructive exhaustion of colloblast adhesive release. We find that colloblast-mediated adhesion is rapid, and that the bonding process is robust against shifts in ambient pH. However, we find that the *Pleurobrachia* colloblast adhesive system is among the weakest biological adhesive systems yet described. We place this surprising observation into a broader ecophysiological context by modeling prey capture for prey of a range of sizes. We find that limited use of colloblast adhesive with high surface area contact is suitable both for capturing appropriately sized prey and rejecting, by detachment, prey above a certain size threshold. This allows *Pleurobrachia*, lacking a mechanism to directly “see” potential prey they are interacting with, to invest in capturing only prey of an appropriate size, decreasing the risk of injury.

**Summary statement:** Ctenophore colloblast adhesive is found to be strong, but few colloblasts are simultaneously active, producing a weakly-adhering system. A physical model demonstrates how such a system may filter unsuitable prey.

## Introduction

Ctenophores are a phylum of gelatinous zooplankton known for being voracious ambush predators (1-4). Most species in the phylum fall within the class Tentaculata (3), characterised by the presence of tentacles whose surface is covered with the phylum-defining colloblast cells (3-7). Colloblasts are bouquet-shaped with an apical enlargement protruding from the tentacle surface. The present study focuses on *Pleurobrachia bachei* and *Pleurobrachia pileus*, two closely related ctenophores of the order Cydippida found in the inshore waters of the Pacific and Atlantic oceans, respectively. Members of this order possess a pair of tentacles with numerous smaller, evenly spaced, tentilla extending perpendicularly off the main tentacle. While hunting, cydippids tend to keep their tentacles and tentilla extended, waiting for prey to drift into the resulting dragnet (3,4). When prey make contact with the tentacles, the colloblasts release their adhesive (likely from a collection of internal vesicles) and bond to the prey (5,6). This discharge presumably destroys the colloblasts, and they may be continually replaced by differentiation from epithelial stem cells (5,8,9).

Ctenophores’ hunting technique is reminiscent of the ambush strategy of orb weaver spiders (2) and the release of an adhesive from colloblasts on prey-contact is superficially similar to the harpoon-like stinging cells in cnidarians, called nematocytes or cnidocytes (4,10-12). However, cnidocytes and colloblasts do not share a common evolutionary origin (13). Many basic questions about colloblasts remain open: What do adhesion events look like? How strong is colloblast adhesive? Understanding the answers to these questions would aid our understanding of colloblast adhesive as a unique biomaterial and inform the potential limitations it puts on ctenophore predation.

In this study, we apply live microscopy techniques to visualise adhesion between probes and tentacles of *Pleurobrachia pileus*, assessing the fates of individual colloblasts engaging in adhesion. We use this system assess the impact of contact area and surface chemistry on adhesion. We then adapt instrumentation for measuring surface tension to measure the adhesive force exerted by the colloblast adhesive system of *Pleurobrachia bachei*. Our data demonstrate that ctenophore prey capture is a robust mechanism that acts quickly to ensnare prey under a variety of conditions. Furthermore, the burgeoning understanding of colloblasts is itself integral to our understanding of ctenophore ecology in a rapidly changing marine environment.

## Methods

### Sample collection

*Pleurobrachia pileus* were collected from the Irish Sea at Howth Harbor, Dublin, Ireland (53.393060, -6.066148) using plankton nets, then transferred into Kreisel tanks established at University College Dublin, School of Medicine. Tanks were circulated with natural sea water (Seahorse Aquariums, Dublin, Ireland) maintained at 10-15 °C, and animals were fed a 20 ml aliquot of live *Artemia* sp. continuously until use within 2 months.

*Pleurobrachia bachei* were collected by dip cupping at the docks at the University of Washington’s Friday Harbor Labs in Friday Harbor, WA. (48.545234, -123.012020). Live animals were shipped overnight to the University of Pennsylvania for testing, where they were held at 4-10 °C in a filtered seawater (FSW) refugium containing macroalgae and live rock from Friday Harbor for experimentation within one week.

### Imaging of Mechanical Probing of Colloblasts

Ecoflex (00-10 hardness) and Dragonskin (10A hardness) silicones (Smooth-On, Macungie, PA) were used to fabricate arenas for probing visualisation. The arenas were placed onto charged microscope slides (Superfrost Plus, Thermo Scientific), and pressed firmly to ensure adhesion to the glass. Anchoring slits were cut into the two corners of the arena. A single portion of severed *P. pileus* tentacle was transferred into the arena with <500 *µl* of natural sea water and orientated using fine-tip tools to anchor the tentacle ends into the anchoring slits.

Probes were generated from 1.75mm diameter filaments of polylactic acid polymer (PLA; Prusa Research, Prague) by heating segments under tension to create tapered strands, subsequently severed to create a flat circular tip. Some probes were immersed in 50 *mg · ml*^*−*1^ poly-D-lysine in PBS until the solution evaporated to coat the tips in lysine. Some probes were adhered to a copepod carcass (*Calanoida* sp. Seahorse Aquariums, Dublin) using glue. Probes were brought into contact with the tentacle fragment (20 *mm · min*^*−*1^), left briefly, and retracted (5 *mm · min*^*−*1^) under 20X magnification video microscopy using an open source microscopy system (14) (Fig. S1).

Probing videos were analysed in ImageJ (15). The proportion of colloblasts actively adhered to the probe in a trial was computed as follows: For any area compressed under the probe (*A*_*compression*_), there will be a certain number of colloblasts contacted. *Pleurobrachia* sp. colloblasts can be represented as circles (∼5 *µm* diameter, 19.63 *µm*^2^ area) densely packed on the surface of the tentacle. Modeling using hexagonal packing arrangement results in a packing density of 0.9, meaning an overall colloblast surface area of 21.63 *µm*^2^ (*SA*_*colloblast*_) when accounting for packing density.

Colloblast activity fraction (CAF) represents the proportion of colloblasts adhered in a probing event. A constant term, B, accounts for any additional factors, such as a minimum number of colloblasts activated in response to any contact (*B >* 0) or a depressive effect within tentacles preventing any colloblast adhesive release until a compression area threshold is passed (*B <* 0). The equation then to calculate the number of colloblasts adhered (*C*_*adhered*_) is,

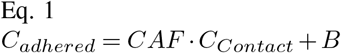

As *C*_*contact*_ is a factor of A_*compression*_ and *Sa*_*colloblast*_, we an rearrange Eq. 1 into the following linear model, where *m* = *CAF/SA*_*colloblast*_

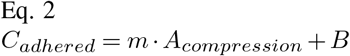

### Assessment of adhesive force and adhesive ability

Measurement of the adhesive force between a probe and a severed tentacle was assessed using *Pleurobrachia bachei*. Tentacles were dissected and immediately laid on clean, untreated glass slides (Corning Inc.; Corning, NY). Excess water was blotted from the tentacles using a Kimwipe (Kimberly-Clark; Roswell, GA), and 200 *µl* of 0.22 *µm* vacuum-filtered seawater was added back to the tentacle to form a droplet around it (Fig. S1). This prepared slide was then positioned in a Kibron EZ-Piplus single channel surface tensiometer with a 0.5 mm diameter DyneProbe probe with a flat contact surface profile and hydrophilic surface chemistry attached (Kibron Inc.; Helsinki). The probe was lowered onto the tentacle at a rate of 0.0125 *mm · s*^*−*1^, and allowed to rest on the surface of the tentacle for a range of time intervals before being raised up, again at 0.0125 *mm · s*^*−*1^. Reported values for adhesive force (*F*_*adhesion*_) represent the difference between the maximum force experienced by the probe as it was being lifted off the tentacle and the force on the probe at the manually determined point of tentacle contact (Fig. S1). Unless otherwise noted, different, randomly selected spots on fresh tentacles were used for each trial, the probe rested on the tentacle for 60 seconds, and the probe was cleaned between trials by heating the tip of the probe until red hot, then wiping the probe down with an ethanol-soaked Kimwipe. In order to assess adhesive depletion/buildup on a probe tip, the adhesive force assay described above was repeated, but modified to probe the same spot on a tentacle multiple times, either cleaning the probe in between these trials (adhesive depletion) or not (adhesive buildup). Two different tentacles were used for these two series of repeated measurements. Assessment of the impact of contact time on adhesive force was performed using the adhesive force assay as described above, but the time the probe was allowed to rest on the tentacle for was varied between 5 and 600 seconds.

To assess the effect of ambient pH on adhesive force, the assay described above was repeated with artificial seawater (ASW) of varying pH. ASW was composed of Instant Ocean Sea Salt (Instant Ocean; Blacksburg, VA) mixed with deionised water to a specific gravity of 1.025. ASW was filtered to 0.22 *µm* and the pH adjusted to the test value using either concentrated HCl or 10 M NaOH as measured with an Accumet AP125 pH meter (Fisher Scientific; Pittsburgh, PA). After mounting the tentacle on the glass slide and blotting away excess seawater, 10 µl of pH adjusted ASW was added to the tentacle to both rinse it and avoid shocking the tentacle with a sudden shift in pH. Then, this water was also blotted away and 200 *µl* of the same solution of pH adjusted ASW was added to form a droplet around the tentacle. This prepared slide was then tested as above. For ambient pH trials, five contacts were made on different locations of three tentacles in each pH condition.

## Results

### Compressive force is a factor in stimulating adhesive release

Direct contact between a PLA probe and severed tentacular surface was not sufficient to trigger colloblast adhesion, indicating a minimum compressive force requirement for adhesive release (Fig. S2A). Retraction of a probe following tentacular compression allowed for visualisation of individual colloblast adhesion events (Fig S2B). Individual colloblast adhesion events were terminated in one of two ways: 1) adhesive failure (de-adhesion event) (Fig. S2B), or 2) colloblast tail severing (Fig. S2C). For probing studies using a soft ridge substrate, no significant correlation between contact area and colloblast adhesion events was observed (Spearman r, *P* = 0.1473, two-tailed, *y* = 0.0000287345*x −*6.88).

When repeating the experiment using Dragonskin silicone as a firm ridge substrate, a significant positive correlation was identified between contact area and colloblast adhesion events (Spearman r, P=0.0467, two-tailed, *y* = 0.000529642*x−*5.474705) (Fig. 1A). However, only a very small proportion of colloblasts (CAF = 1.04%) are activated in response to compression under these conditions.

**Figure 1.**
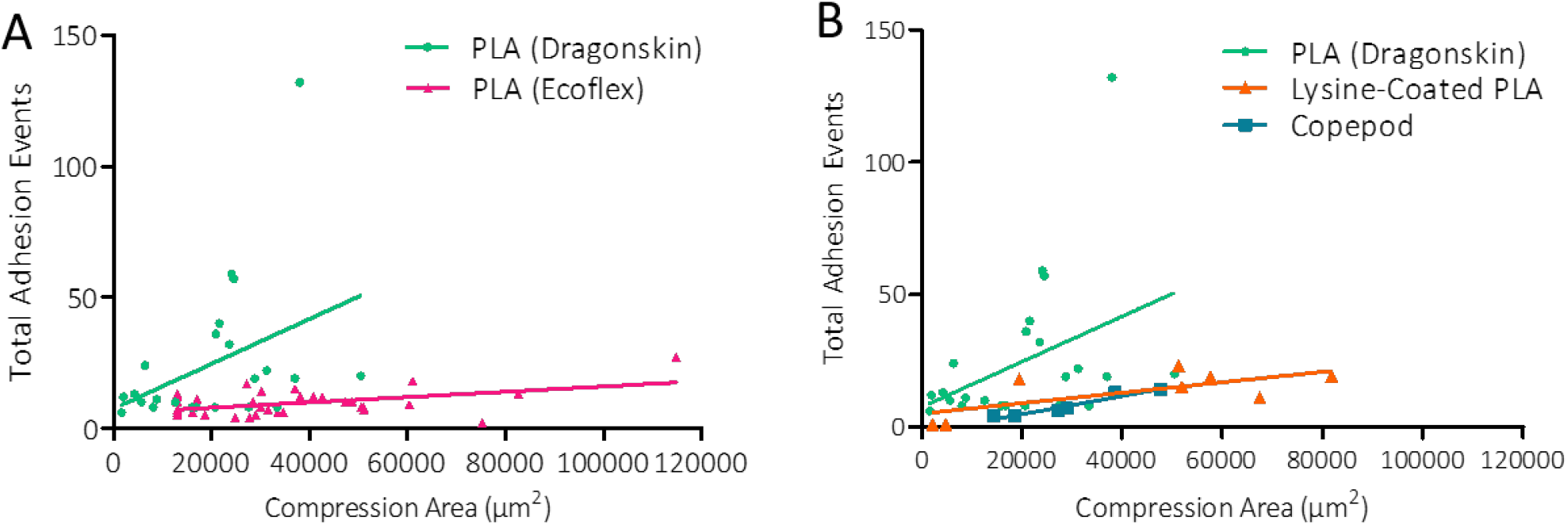
Parameters of Probe Compression Impact Colloblast Adhesion Events. A) Tentacle compression against firm Dragonskin resulted in a significant positive correlation between compression area and total adhesion events (Spearman r, P=0.0467, *y* = 0.000529642*x* + 5.74706) but not against Ecoflex (Spearman r, P=0.1473, *y* = 0.0000287345*x* + 6.88). B) Total adhesion events were plotted against the area of compression induced by the probe. Significant positive correlation was also identified using copepod carcass in a Dragonskin arena (Spearman r, P=0.0028, *y* = 0.00025*x*4.35411), but not in lysine-coated probes (Spearman r, P=0.1966, *y* = 0.00013*x* + 23.88775). All mathematical lines of fit (LOF) were acquired using Theil’s incomplete method for non-parametric data, however visual representation of LOF is calculated without compensating for non-normal distribution (Prism).

### Effect of probe surface chemistry on adhesive response

Repeating probing experiments using probes coated in hydrophilic lysine revealed no significant correlation between probed area and the number of adhesion events (Spearman r, P=0.1966, two-tailed), however due to the low number of experimental repeats (n=8) further studies are warranted. A significant correlation between probed area and the number of adhesion events was observed when probing using a copepod carcass (Spearman r, P=0.0028, two-tailed, *y* = 0.00025*x −*4.35411), however the proportion of colloblasts activated in response to probing (0.491%) was less than half the proportion activated in untreated probe probing. During probe retraction from tentacles, it was possible to visualise the point at which deadhesion events occurred, these distances were collated for all untreated probe contact events (Fig. 2C), for which the median deadhesion event distance was 83.33 *µm*. This was significantly greater than the median deadhesion event distance for lysine-coated probes (16.48 *µm*, Dunn’s Multiple Comparison test, P<0.0001) or copepod probes (10.73 *µm*, Dunn’s Multiple Comparison test, P<0.0001). However, the distance of deadhesion events was not found to be significantly different between lysine-coated probes and copepod carcass probing (Mann Whitney, P=0.7972).

**Figure 2.**
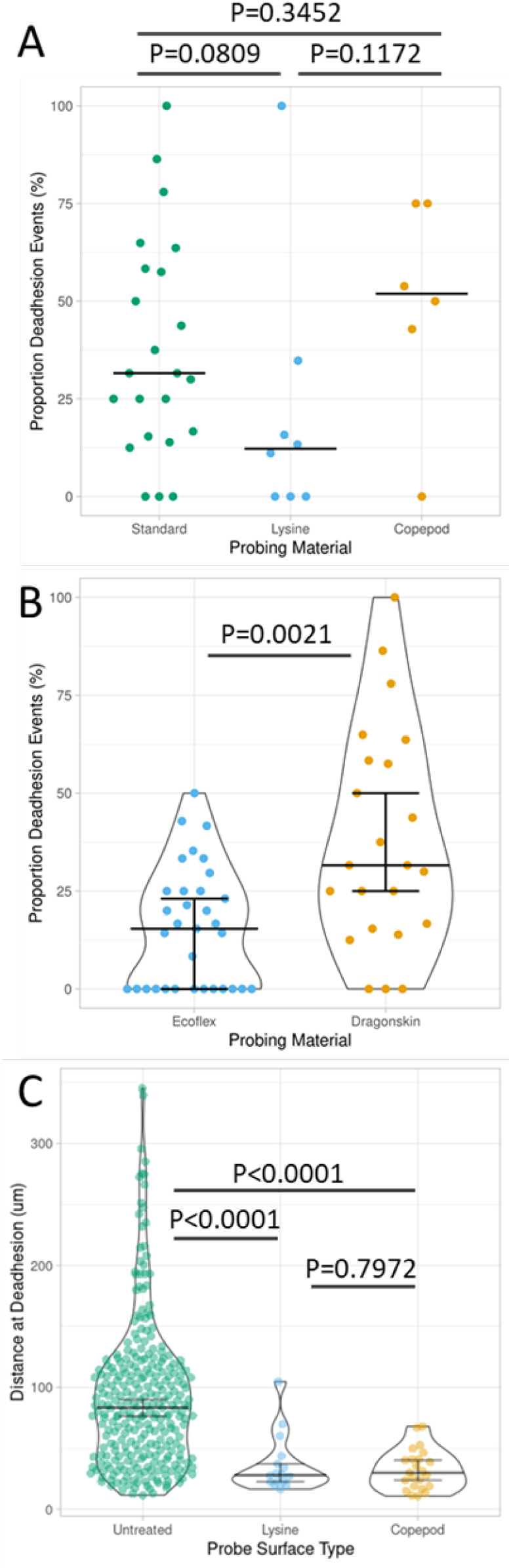
Surface Chemistry of the Probing Surface Impacts Distance of Deadhesion, but not Termination Modality. A) Proportion of observed adhesion termination events classified as deadhesion events for each probe type used using Dragonskin arenas. Median shown as solid black bar. B) Proportion of observed adhesion termination events classified as deadhesion events using untreated PLA probes with either an Ecoflex or a Dragonskin ridge. Bars show mean +/-95% CI. Statistics performed using Mann Whitney test. C) Distance (µm) between tentacle surface and adhered colloblast head at time of deadhesion, compared between probing using untreated PLA probes, lysine-coated PLA probes, and copepod carcasses, statistical analyses performed using Kruskal-Wallace and Mann Whitney tests.

### Compressive force and observed deadhesion modalities

The proportion of deadhesion and colloblast tail severing adhesive termination events were compared between the probing types using Dragonskin as a ridge substrate, and this proportion was not found to be significantly different (Kruskal-Wallis test, P = 0.1136, *n*_*P LA*_ = 23, *n*_*lysine*_ = 8, *n*_*copepod*_ = 6), indicating the mode of adhesive termination is not affected by probing surface chemistry (Fig. 2A). However, the proportion of deadhesion events was found to differ significantly between untreated PLA probing with an Ecoflex ridge vs a Dragonskin ridge (two-tailed Mann Whitney test, P = 0.0021), with Ecoflex resulting in a significantly lower proportion of deadhesion events compared to Dragonskin (Fig. 2B), indicating compressive force plays a role in the modality of adhesion termination events.

### Baseline Force and Adhesive Buildup/Depletion Assays

We conducted two series of repeated measurements of the adhesive force arising from a single spot on two different *P. bachei* tentacles (Fig. 3). In one series, we cleaned the measuring probe between measurements, removing any adhesive substances released by the colloblasts. In the other series, the probe was not cleaned, allowing any adhesive substances released by the colloblasts to build up on the probe’s surface. Both series began with a starting value of about 200 micronewtons (*µN*) of adhesive force, which we subsequently take as a rough “baseline” adhesive force value. In the series of trials with probe cleaning, the adhesive force drops asymptotically to zero (fit to the model: *y* = 386.8 *e*^*−*0.6781*x*^ with *R*^2^=0.99). In the series of trials without probe cleaning, the measured adhesive force hovered around the baseline value of 200 *µN* for the first three trials, after which it gradually increased before plateauing around 300 *µN*.

**Figure 3.**
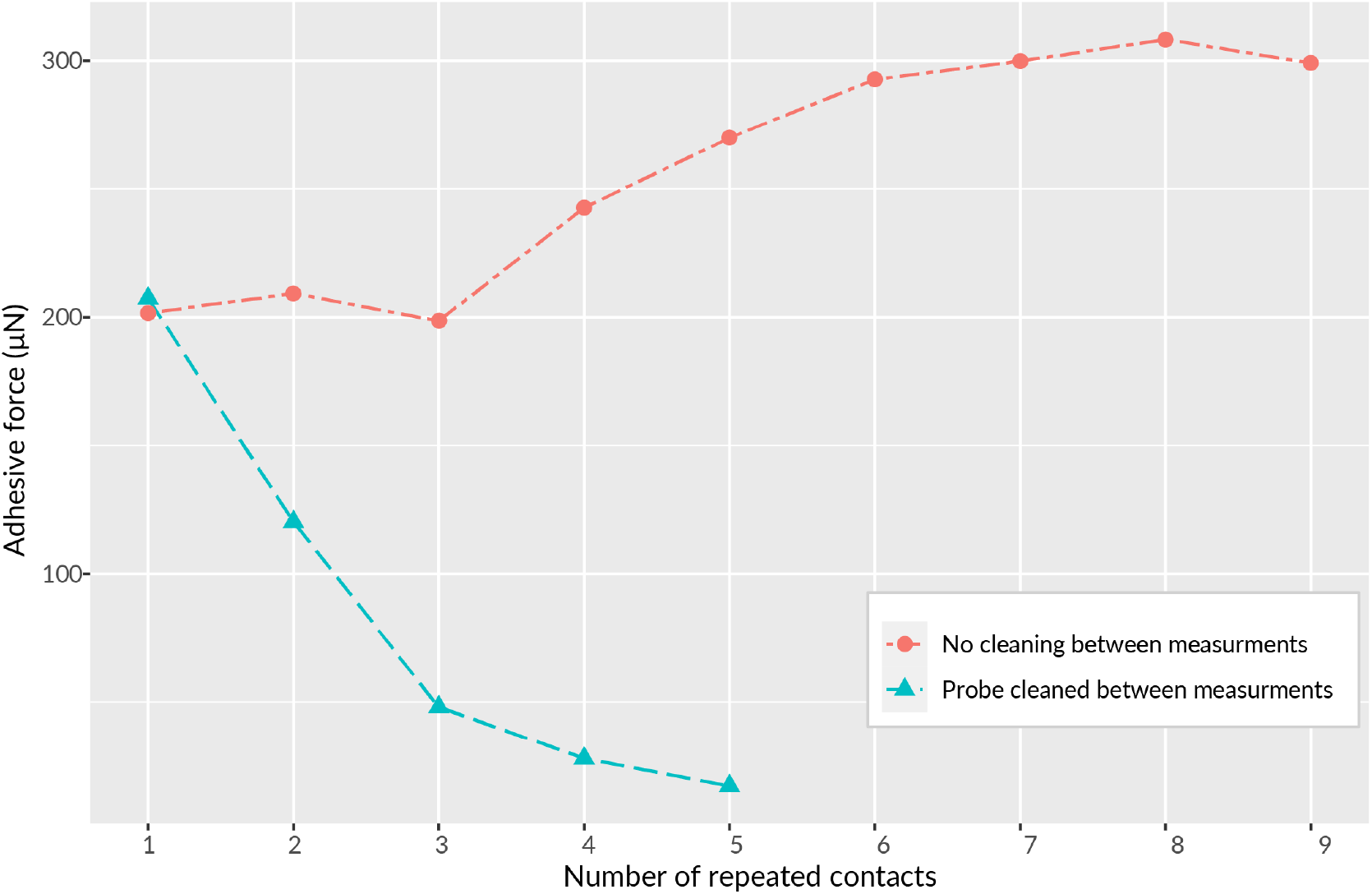
Plot of adhesive forces from repeated measurements at a single spot on two different *P. bachei* tentacles with two different testing protocols. Between trials, the testing probe was either cleaned, presumably removing any residual adhesive, or left uncleaned, presumably allowing adhesive to build up on the surface of the probe until all available adhesive at that spot is depleted.

### The effect of curing time on adhesive force

Though cydippid ctenophores ambush and ensnare their prey, and adhesion is thus presumed to happen rapidly, it is unclear precisely how rapidly adhesion may occur or if the bond may increase in strength if allowed to cure for longer amounts of time. To investigate this, we assayed the effect of curing time on colloblast adhesion by probing the tentacle, this time at a fresh spot on a new tentacle between trials, while varying the length of time the probe was allowed to sit and “cure” on the tentacle surface for either 5, 60, 180, or 600 seconds (Fig. 4). The lowest curing time value of 5 seconds was limited by the response time of the testing apparatus. We observed no statistically significant difference in bond strengths between different curing times (Fig 6.), as assessed by Kruskal-Wallis testing (P-value cutoff of 0.05). During adhesion visualisation experiments, several instances of adhesion were observed following compression for a much shorter duration than 5 seconds. In some instances, adhesion occurred instantaneously with compression, visualised as colloblast adhesion events as the tentacle contracted prior to probe retraction.

**Figure 4.**
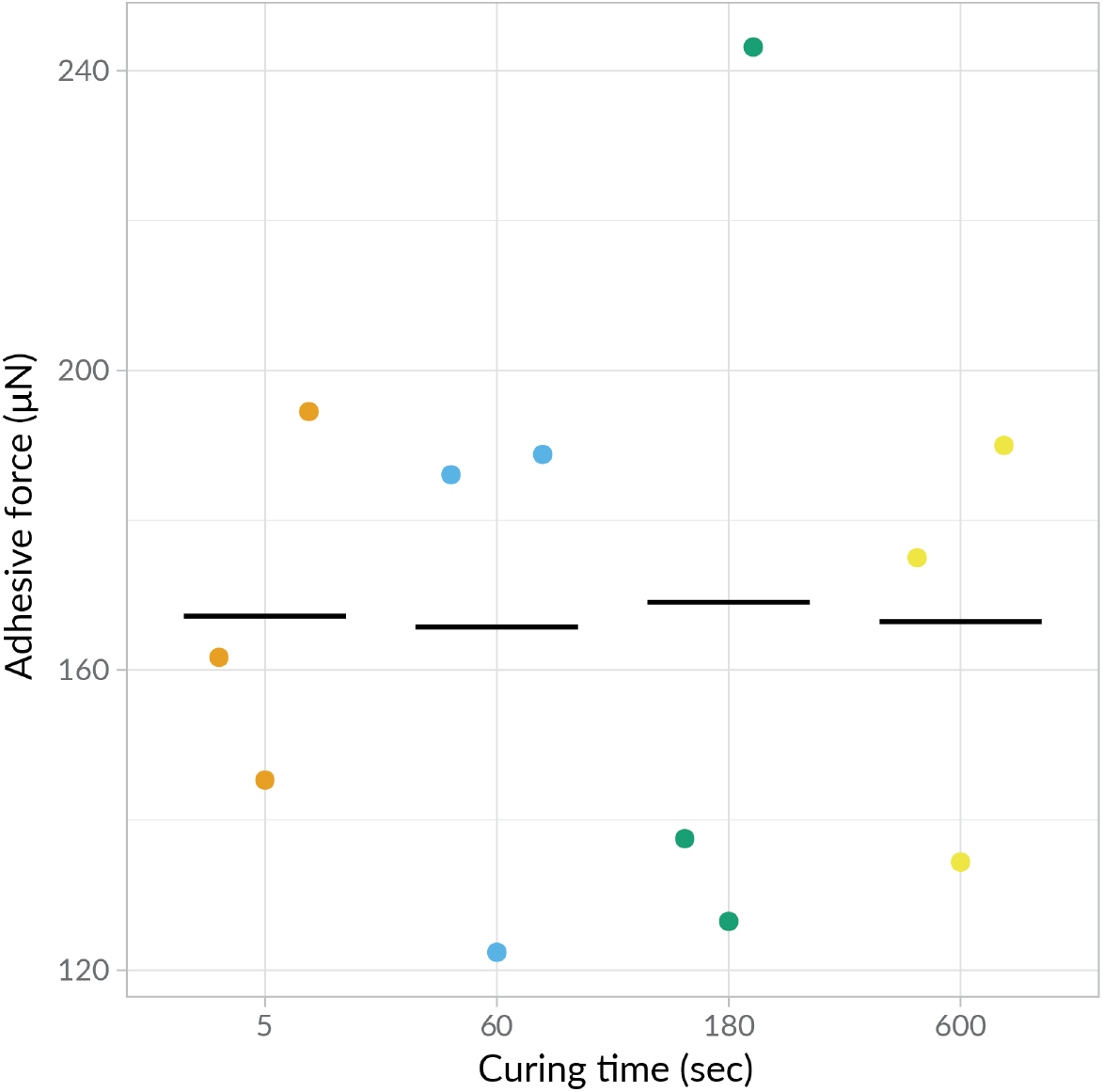
Adhesive force did not significantly depend on the amount of time that the adhesive was allowed to cure. No curing time trial groups’ medians differ by Kruskal-Wallis rank sum test, suggesting that tP. bachei colloblast adhesive both acts rapidly and once attached, may remain bonded for some time. Filled circles = data points; black lines = mean values.

**Figure 5.**
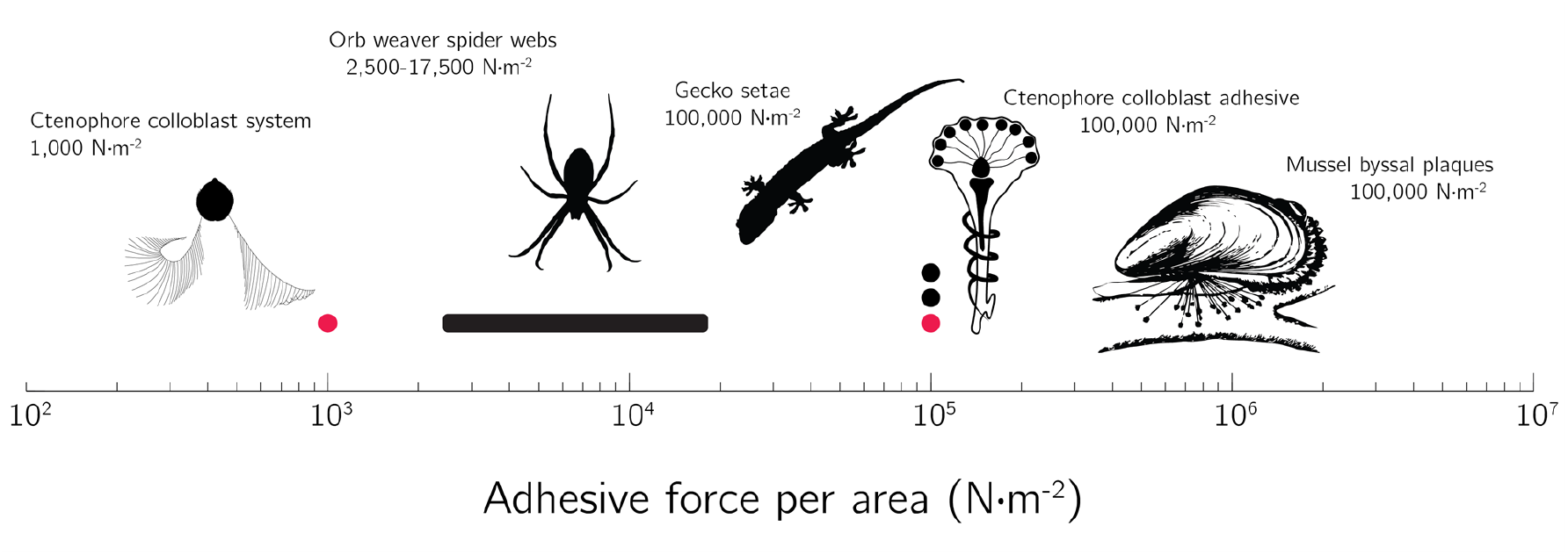
Comparison of various biological adhesive systems by adhesive force per unit area. The adhesive force generated by the colloblast adhesive system is among the lowest of biological systems that have been examined, though the adhesive itself is relatively strong. Ctenophore, orb weaver spider, and gecko silhouettes downloaded from the PhyloPic database (http://phylopic.org). Mussel silhouette created from a public domain image from the Freshwater and Marine Image Bank at the University of Washington. Pink circles, force values observed in *Pleurobrachia*; black circles, forces observed in non-ctenophore adhesive systems.

**Figure 6.**
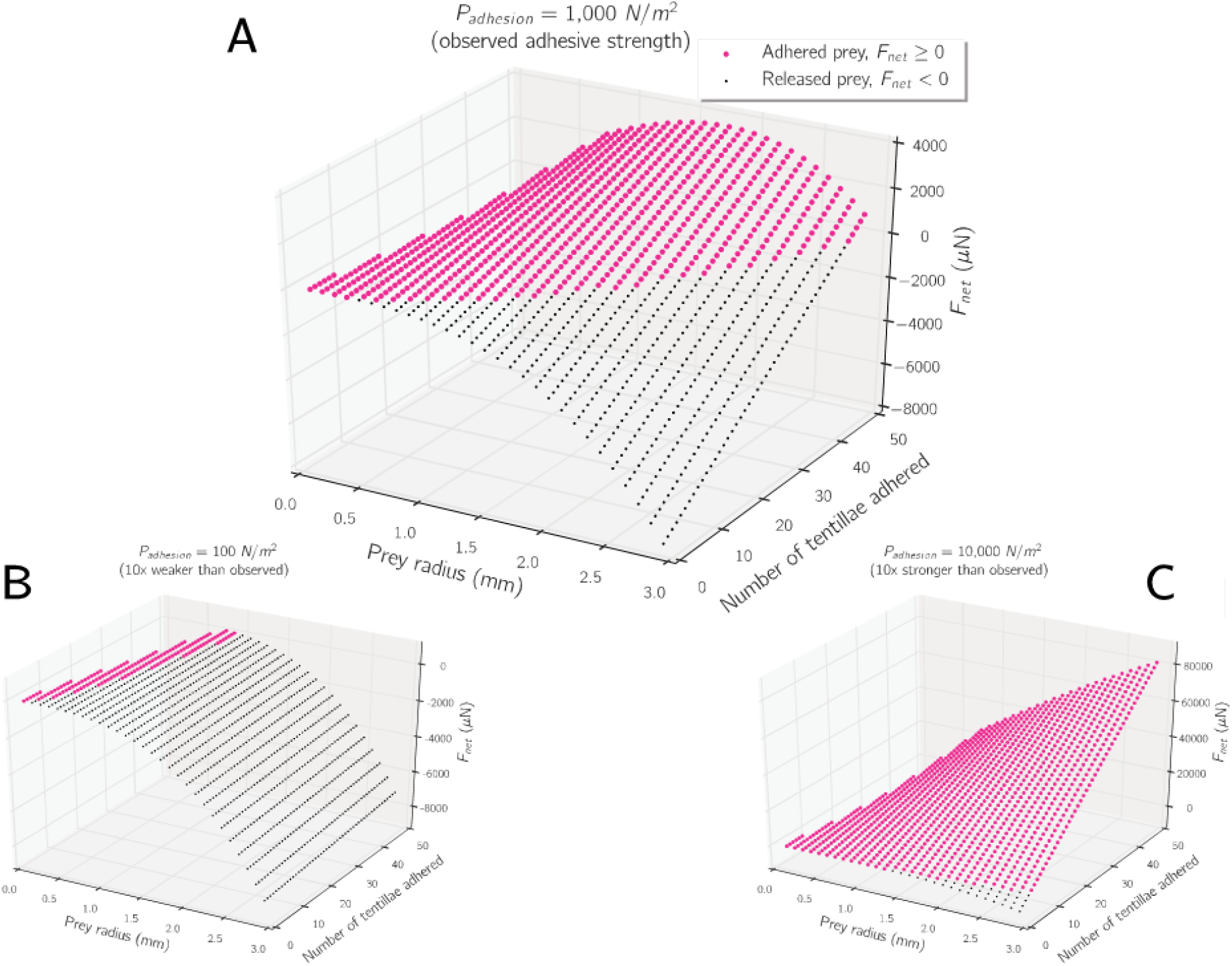
Modelling the net force on hypothetical escaping prey items adhered to multiple ctenophore tentilla. If *F*_*net*_ is positive, the adhesive force exceeds the opposing escape force, and the prey remains stuck (pink circles). If *F*_*net*_ is negative, the escape force wins out, and the prey is released from the tentilla (black circles). Only an adhesive of the approximate order of magnitude strength as we observed (A) allows for a relatively wide range of acceptable prey while still allowing for the release of very large prey (1.5 mm radius). If the adhesive’s strength were 10 times weaker (B), only a very narrow range of very small prey could be captured, whereas if it were 10 times stronger (C), prey items of unmanageably large sizes not reliably handled in nature would easily adhere to even a single tentillum, potentially exposing the animal to injury.

### Impact of ambient pH on colloblast adhesive force

Ambient pH can significantly affect aquatic life in general and chemical adhesion more specifically, an effect that has been studied extensively in the adhesive system of mussel holdfasts (16–18). With this in mind, we measured the adhesive force of *P. bachei* tentacles in artificial seawater adjusted to four different pH conditions: 9.0, 8.2 (approximately “standard” ocean surface pH), 7.0, and 6.0 (Fig. S3). Kruskal-Wallis testing showed a significant difference between the different test conditions (P=0.04) and subsequent pairwise Dunn’s testing with Bonferroni correction showed that the only statistically significant difference between the medians was that between pH conditions 7.0 and 8.2 (P=0.0138). We also calculated the moment coefficient of skewness for each condition, and found that the data for pH conditions 9, 7, and 6 all have positive skew coefficients (0.40, 0.56, and 0.81 respectively), with mean values exceeding the median values in each of these conditions. This is due to prominent tailing of the data toward higher values of adhesive force, visible in the kernel density estimates around these data in Fig. S3.

## Discussion

### A subset of colloblasts release adhesive when stimulated

The results presented here support three commonly presumed properties of colloblast adhesion —the adhesive is an expendable substance released from the colloblast, the adhesive is fast-acting, and the adhesive can be released in response to physical stimulation (4–6). However, visualisation of tentacle probing showed that adhesive is released from only a small proportion of colloblasts. Repeated probing of areas in our force measurement studies showed increasing adhesion strengths up to a plateau and are consistent with a system in which colloblast activation and adhesive release are stochastic events, with each serial probing event triggering a proportion of the remaining colloblasts to release adhesive. When the probe is cleaned between trials, this appears as a depletion of adhesive force in that spot, and when the probe is not cleaned between trials, this appears as adhesive building up on the probe, gradually reaching a maximum (300 µN). Based on our estimates of colloblast density, the maximum expected number of colloblasts in the area probed would be 9,000, and based on our maximum estimate of colloblast activation we would expect only 100 to be activated and release adhesive (based on PLA on Dragonskin). Thus, the force measurement maximum would be expected to be 2 orders of magnitude higher than that recorded if accumulation of the total adhesive was occurring. While it was not possible to visualise repeat probing of a single area, we have visualised adhesive and colloblast heads deposited on the surface of a probe tip following retraction of a probe, potentially preventing adhesive from re-adhering to the tentacle on re-probing. Adhesive curing occurs faster than re-probing can occur, and thus the originally deposited adhesive may no longer be sticky by the second round of probing. An alternative explanation could be that repeat probing of tentacles does not trigger all colloblasts to eventually be triggered, representing desensitisation of the adhesive release system and potentially adding to evidence of a common neural origin of colloblast cells (13). The lack of adhesion observed with a light tapping of tentacles indicates some threshold of physical force required for adhesive release, which is corroborated by the total adhesion event number findings when comparing soft with firm arenas during probing visualisation. A significant difference between the modality of adhesion termination events implicates force imparted as a modulator of adhesivity of colloblasts. Counterintuitively, lower compressive forces applied would seem to indicate a stronger adhesive bond between the colloblasts and the probes.

### Colloblast adhesive cures rapidly under variable environmental conditions

The adhesive in ctenophore colloblasts also acts rapidly: the adhesive force measured when the probe had been allowed to cure for 5 seconds was comparable to the force measured when it had been allowed to cure for 10 minutes. The adhesive is likely fully cured in less than 5 seconds’ time, as shown in incidental recordings of low contact-time probing during the visualisation studies. Given that tentaculate ctenophores are ambush/entanglement-predators not unlike spiders in their approach to feeding, this is a reasonable conjecture.

Interestingly, colloblast adhesive appears to function with similar efficiency across the wide range of ambient pH values tested. This implies that either the adhesive substance and its chemical mechanism of adhesion are relatively resistant to shifts in pH or that the pH of the adhesive granules themselves is controlled or buffered and adhesion occurs rapidly after the granules rupture, leaving little time between granule rupture and bonding to prey for pH to effect adhesion.

### Colloblast adhesive is relatively strong, but the colloblast adhesive system is relatively weak

Force measurement utilised a tensiometer probe with hydrophilic surface chemistry. This type of surface is useful because it strongly maintains a layer of water around the probe that must be displaced or coordinated with for colloblast adhesive to adhere to the probe’s surface. Ctenophore prey items in the wild likely run a wide gamut of surface chemistries that are difficult to characterise in practice, but our approach provides a reasonable baseline by making water a significant barrier to interacting with the probe surface.

If we compare the adhesive strength of *Pleurobrachia* colloblasts to other studied biological adhesive systems in terms of force per unit area, colloblast adhesive appears relatively weak (Fig. 5). Based on Eq. 2 the expected number of colloblasts initially adhering to the force measurement probe is 100 colloblasts, yielding an estimate of ∼2 µN of adhesive strength per active colloblast (1,000 *N · m*^*−*2^). If all 9,000 colloblasts contacted by the probe activated at once, this would result in ∼18,000 *µN* of adhesion across ∼0.2 *mm*^2^, or 90,000 *N · m*^*−*2^, a rough estimate of the strength of the adhesive considered on its own.

This means that the strength of colloblast adhesive itself is on par with that of systems involved in anchoring whole animals such as mussel holdfasts (Fig. 5), and gecko foot setae (19–21). This is particularly notable given that mussel holdfast adhesives may be biochemically similar to Pleurobrachia colloblast adhesive (22). However, the adhesive strength of the colloblast system as a whole is much weaker than these whole-animal anchoring systems, and under working conditions colloblasts adhere to prey with a force on the same order of magnitude as orb weaver spider webs’ viscous capture threads (23).

### Modelling of ctenophore prey capture with a weak adhesive system suggests a mechanism for prey selection

It’s not immediately obvious why *Pleurobrachia* might employ a strategy of using a strong adhesive sparingly to produce a weakly-adhering system. Our data allow us to suggest one potential explanation for this strategy. Here, we present a model describing how a relatively weak adhesive system can mechanically filter prey by size.

*Pleurobrachia* commonly prey upon small animals such as copepods (3,4). They utilise an ambush hunting strategy reminiscent of orb-weaver spiders (2) —extending their tentacles and fanning out their numerous secondary tentilla, waiting for prey to close in, and ensnaring them in this sticky “web”, with closely arranged tentilla increasing the chance of prey contact with multiple tentilla. As a biological apparatus, damage to the tentacles/tentilla represents a risk to the animal, and so the need to capture prey must be balanced with the need to protect the animal’s prey-capture ability.

Consider a hypothetical prey, modeled as a sphere with a radius, *r*_*prey*_ that has values within a range of 0.1–3.0 mm, which encompasses the range of nauplii and copepod sizes (radii of approximately 0.11–1.5 mm) for which data on *Pleurobrachia* prey capture exist in addition to hypothetical prey larger than those known to be reliably captured by *Pleurobrachia*. This prey has a mass, *m*_*prey*_ based on a copepod density value, *ρ*_*prey*_, of 1,050 *kg · m*^*−*3^ as estimated from available literature (24), thus,

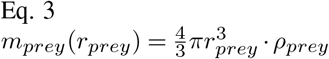

When an individual tentillum, measured from our micrographs to be 30 microns wide (*w*_*tentacle*_), makes contact with the prey, it wraps a third of the way around the sphere’s circumference and adheres with a contact area, *A*_*contact*_

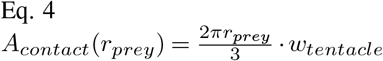

In this model, additional tentilla can contact the prey and potentially adhere in the same fashion. A key limitation of our visualisation studies is that they relied on linear segments of tentacle probing, along with a probing force applied in a single direction, contrasted with a natural setting in which multiple tentilla could be contacted by a prey which would be moving vigorously with unpredictable direction. The total adhesive force is proportional to the surface area of all tentilla adhered to the prey, given the measured adhesive strength value, *P*_*adhesion*_, of ∼1000 *N · m*^*−*2^. As an escape response, prey accelerate directly away from the adhering surface of the tentacle, generating a maximum force proportional to their mass (25). For this model, we consider a representative value for the mass-specific force of 100 *N · m*^*−*2^ (denoted as *a*_*prey*_), allowing us to model total adhesive force and escape force, and also oppose them to plot the difference between these forces

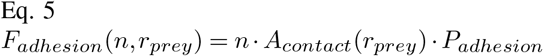

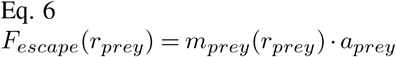

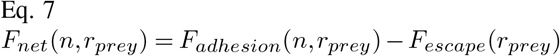

When *F*_*net*_ is positive, the force of adhesion is greater than the force of the prey attempting to escape, and the prey remains adhered. We consider n within a range of 1 (a single tentillum) to 50 (the approximate total number of tentilla on an adult *Pleurobrachia* tentacle). We compute *F*_*net*_ only when the condition *n*.*w*_*tentacle*_ *≤* 2*r*_*prey*_ is satisfied, a rough boundary condition reflecting that a prey item cannot have more than its diameter’s worth of tentilla adhered across its surface. Plotting *F*_*net*_(*n, r*_*prey*_) using the observed value for *P*_*adhesion*_ in *Pleurobrachia* tentacles of 1,000 *N · m*^*−*2^ (Fig. 6A), reveals several interesting patterns. Small prey items with radii less than 0.5 mm adhere and stay adhered on contact with even a single tentillum. Making contact with other tentilla only enhances this effect. A medium-sized prey item with a radius of 0.5-0.7 mm must make contact with two to three tentilla in order to stay adhered, while large prey models (e.g. radius of 1.5 mm) escape unless 12 or more tentilla adhere to it. The largest prey models shown, between 2.5 and 3.0 mm, will not stay adhered unless they make contact with two thirds or more of the tentilla on the entire tentacle. Because *F*_*adhesion*_ is dependent on the available contact area, and thus the prey radius, the slope of *F*_*net*_ as a function of tentacles adhered increases with increasing prey size.

The overall pattern of size-dependent prey capture rates depicted in this model is consistent with laboratory observations of *Pleurobrachia* feeding (2) with approximately 0.5-1.0 mm spacing between tentilla. This makes the capture of larger prey less likely due to the low probability of contact with sufficient tentilla to overcome the escape response. Lacking any visual means to sense the size of prey, ctenophores need some means to accept prey of a suitable size and reject those it cannot due to risk of damage and the metabolic cost of tentacular regeneration (26,27). In our model, prey are mechanically sorted by size via a low strength adhesive system requiring multiple points of contact in order to capture larger prey, minimizing the risk of failed investment in overly large prey. The low proportion of colloblasts activated in prey-contact reduces the risk of losing excessive numbers of colloblasts in the event of a failed feeding event, ensuring the tentacle is capable of re-adhering to new prey in a short timeframe. If we consider the same model as described above, modifying only the value of *P*_*adhesion*_ to be 10 times weaker (100 *N · m*^*−*2^) we obtain a model ctenophore that needs many tentilla to adhere to catch even the smallest prey items (Fig. 6B), and is thus unlikely to capture all but the smallest prey. If the adhesive strength parameter is made 10 times larger than the observed value (*P*_*adhesion*_= 10, 000 *N · m*^*−*2^) we see that most model prey items considered strongly adhere with only one or two tentilla attached (Fig. 6C). Even prey with diameters of 6 mm, a size approaching the centimeter length scale corresponding to the total body length of an adult *Pleurobrachia*, require only 5 tentilla to adhere for capture, presenting increased risks and reduced chance of reward. The measured adhesive system strength of 1,000 *N · m*^*−*2^ allows Pleurobrachia to occupy a biomechanical medium, where a range of reasonable prey sizes can be reliably handled, while larger, potentially dangerous prey are allowed to safely detach.

## Concluding Remarks

We measured the adhesive strength of the colloblast adhesive system in the ctenophore *P. bachei* and demonstrated that colloblast adhesive is released destructively and bonds to its target in seconds. Visualization of probing events using *P. pileus* confirmed this destructive process through visualization of colloblast detachment from tentacles and allowed for the calculation of colloblast adhesive strength. Furthermore, we showed that the bonding of this adhesive was robust to a wide range of ambient pH conditions. Surprisingly, while we found that individual colloblasts are not weak, the adhesive system as a whole is very weak, as the majority of colloblasts do not activate on contact. By considering colloblast adhesive in the context of *Pleurobrachia* feeding behavior and modeling a range of prey size and tentilla contact scenarios, we postulate that this low-strength adhesive system may serve a key purpose. *Pleurobrachia* rely on prey colliding with the animal’s meshwork of tentilla, which are themselves an organ whose damage would put the animal at a disadvantage in feeding. With no way to otherwise “see” directly what prey they are catching or how massive it is, ctenophores need a mechanism to select for appropriate prey items. The colloblast adhesive system’s relatively low strength means that for all but the smallest prey items, multiple points of contact are necessary to trigger sufficient adhesive release to securely handle potential prey, and that above a certain size, it is unlikely that enough contact can be made to secure the catch, allowing these very large prey items to safely terminate adhesion without doing substantial damage to the ctenophore. These observations suggest that the control of ctenophore colloblast adhesive release may be tuned to facilitate the complex task of passively selecting appropriate prey.

## Authors’ contributions

JPT conceived of the study, designed and performed force measurements and data analysis, developed biophysical models, and drafted the manuscript; GOTM designed and performed live microscopy experiments and data analysis, and critically revised the manuscript; GPC performed adhesive force measurements and data analysis; MP designed live microscopy experiments and critically revised the manuscript. All authors gave final approval for publication and agree to be held accountable for the work performed therein.

## Acknowledgements

Drs. Alison Sweeney, Paul Janmey, Mike LaBarbera, Jack Costello, and Thomas C Collin provided comments that improved the manuscript.

## Supplementary Materials

**Figure S1.**
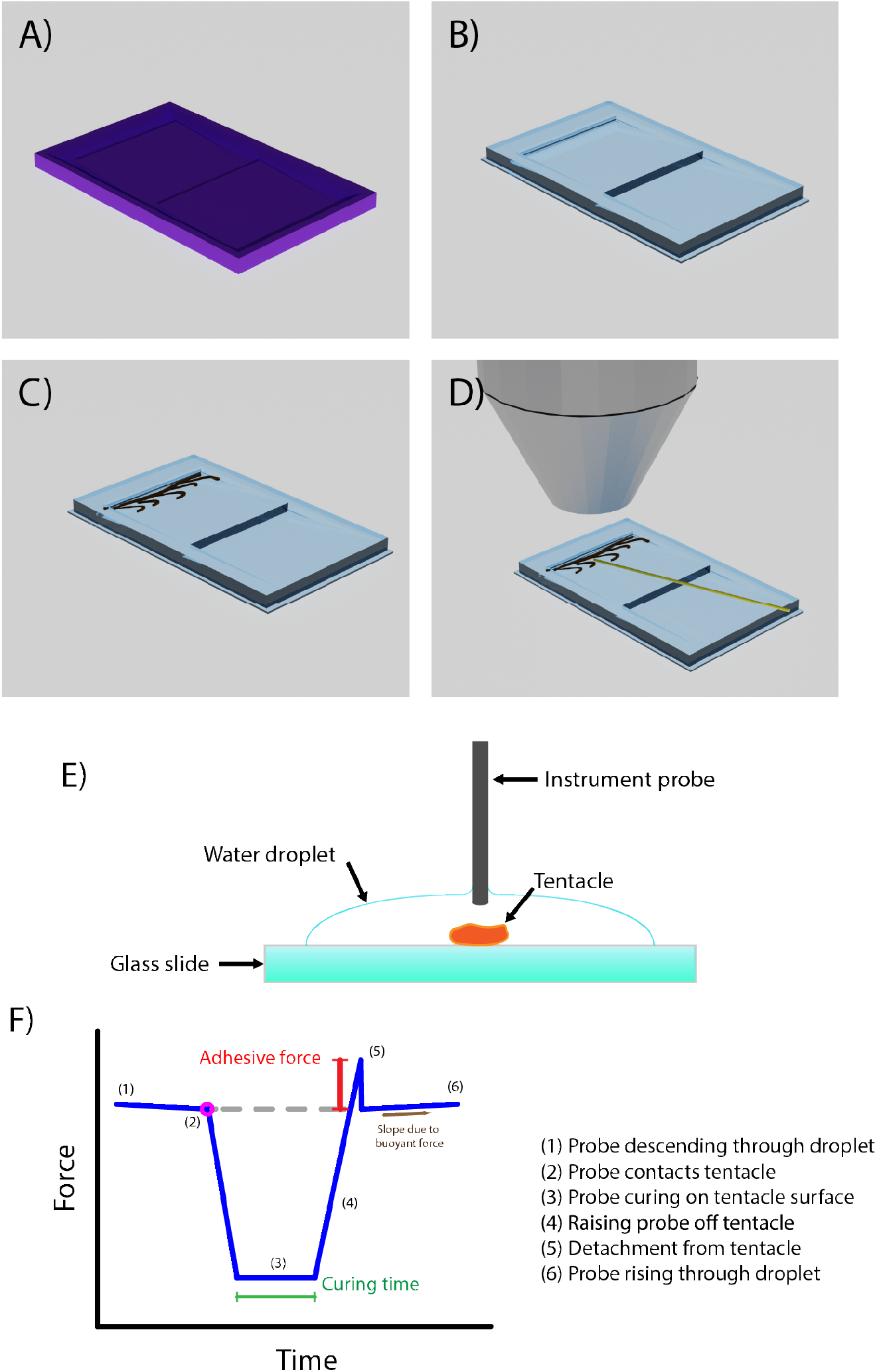
Experimental Methodology Design. Probing visualization: A) A 3D printed mould with two sloped regions was printed in PLA (0.15mm z-resolution, Prusa MK3). B) A negative of this mould was cast using either a firm silicone (Dragonskin shore hardness 10A) or soft silicone (Ecoflex, shore hardness 00-10), cured, and removed from the mould. C) A severed portion of tentacle from *P. pileus* was orientated within the ridge portion of a slope with ∼500 *µl* of natural sea water. D) A probe of either PLA, lysine-coated PLA, or a copepod carcass was orientated using an XYZ micro-translator to be in proximity to the tentacle under 20X objective magnification. The micro-translator was mechanically coupled to the optics. For probing, the stage (which is decoupled from the optics/probe) was moved to bring the tentacle into contact with the probe under oblique video microscopy. Force measurement: E,F) A dissected tentacle is laid on a clean glass slide and surrounded by a droplet of seawater. The experimental probe of a tensiometer is lowered through the water onto the tentacle and allowed to adhere to its surface. As the probe descends through the droplet, it is opposed by a buoyant force proportional to the volume of probe that has been submerged (cf. negative slope of the force trace before the magenta circle). At the point of contact with the tentacle (magenta circle), this gradual decrease in force stops and is replaced by a sudden, sharp drop to approximately no force as the probe lifts off from the tensiometer’s cantilever sensor as the probe can no longer move down. Then, the probe is retracted from the tentacle until it detaches from the tentacle surface. The difference between the force on the probe as it makes context with the tentacle and the maximum force it experiences before detachment is the “adhesive force” we report (red bar with whiskers). It should be noted however, that this value is in fact a pressure, a force experienced over the contact area of the probe with the tentacle.

**Figure S2.**
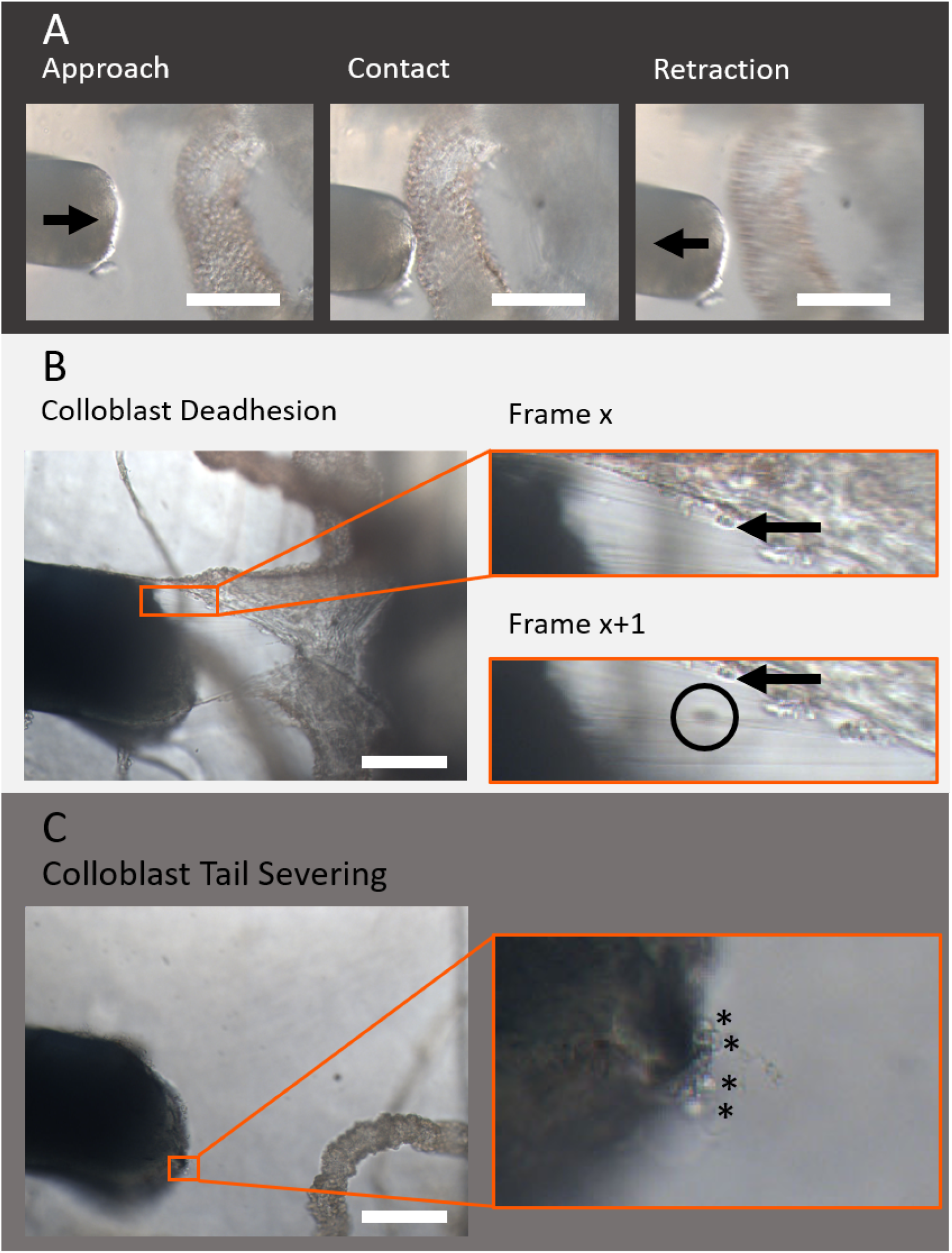
Compression of Tentacles was Required for Adhesion Events and Adhesive Termination to be Visualized. A) Representative images of probe contact without tentacular compression. The probe is brought towards the tentacle until contact is just made. The probe is left stationary for 5 seconds, then retracted. No probing experiments of this type resulted in visualization of adhesion events. B) Representative images highlighting visualization of deadhesion events. As the probe is retracted zooming in to the space between the probe tip and the tentacle core reveals colloblast tails tethering adhered heads on the probe tip to the tentacle core. Further retraction allows for visualization of a colloblast head (black circle) springing back towards the tentacle, having deadhered from the probe tip. Arrow is used to denote landmark feature within both frames. C) Representative images highlighting visualization of colloblast tail severing events by identification of colloblast heads on the probe tip surface following full retraction from the tentacle. The probe is retracted and zoom applied to the probe tip. Colloblast heads appear as spherical objects (*) adhered directly to the probe tip with no colloblast tail tethering it to the tentacle. All scale bars represent 100 *µm*.

**Figure S3.**
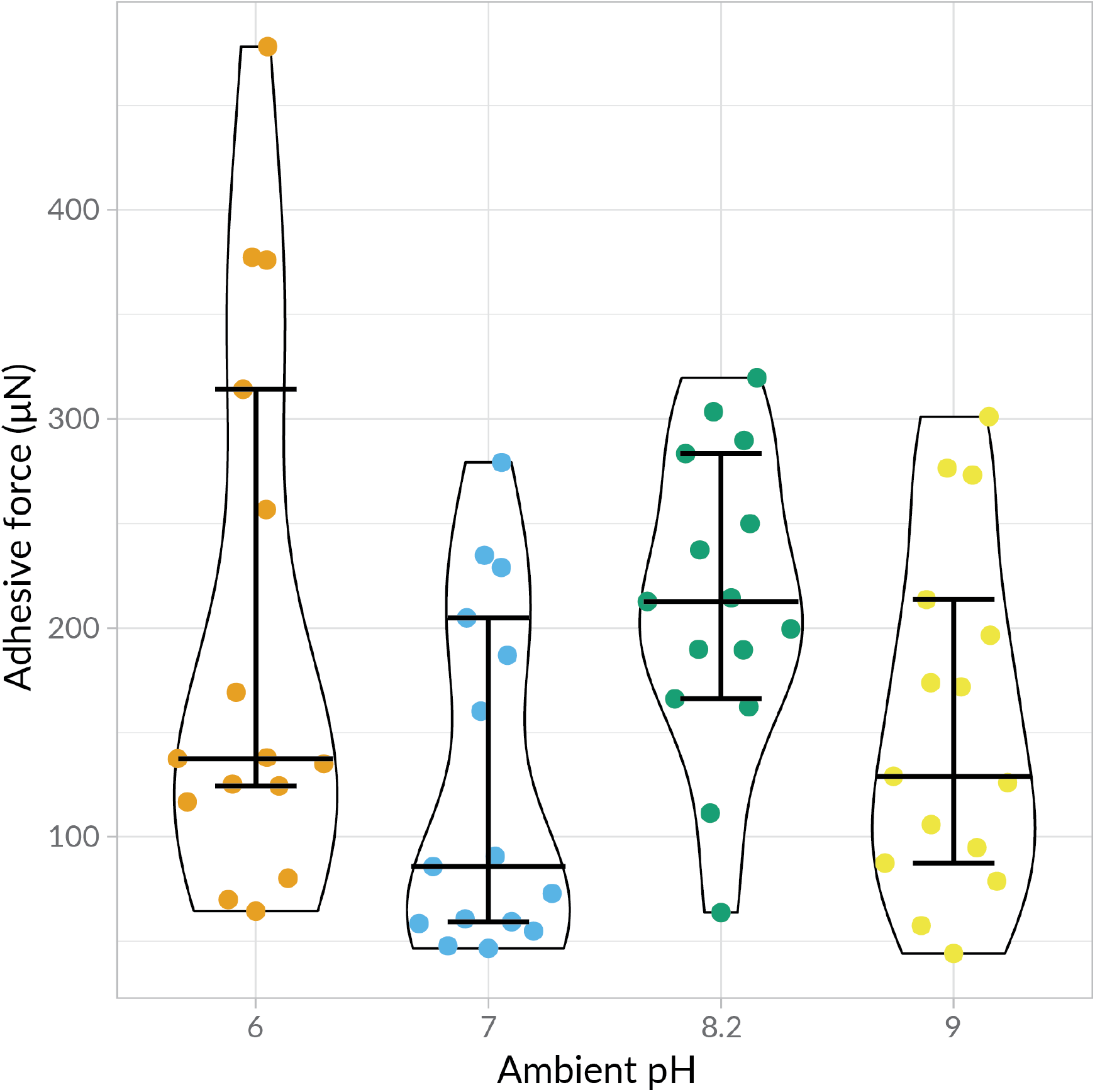
Under varying ambient pH conditions *P. bachei* tentacles show similar median adhesive forces. Kruskal-Wallis testing followed by pairwise Dunn’s tests showed that adhesive strength differed significantly only between the ambient pH 7 and 8.2 trial groups. The distribution of observed adhesive forces in all trial groups besides pH 8.2 showed substantial positive skew, visible in the kernel density estimates of the above violin plots. Black bars = 95% confidence interval about the median..

